# An improved tetracycline-inducible expression system for fission yeast

**DOI:** 10.1101/2024.06.22.600193

**Authors:** Xiao-Hui Lyu, Yu-Sheng Yang, Zhao-Qian Pan, Shao-Kai Ning, Fang Suo, Li-Lin Du

## Abstract

The ability to manipulate gene expression is valuable for elucidating gene function. In the fission yeast *Schizosaccharomyces pombe*, the most widely used regulatable expression system is the *nmt1* promoter and its two attenuated variants. However, these promoters have limitations, including a long lag, incompatibility with rich media, and unsuitability for non-dividing cells. Here, we present a tetracycline-inducible system free of these shortcomings. Our system features the *enotetS* promoter, which achieves a similar induced level and a higher induction ratio compared to the *nmt1* promoter, without exhibiting a lag. Additionally, our system includes four weakened *enotetS* variants, offering an expression range similar to the *nmt1* series promoters but with more intermediate levels. To enhance usability, each promoter is combined with a Tet-repressor-expressing cassette in an integration plasmid. Importantly, our system can be used in non-dividing cells, enabling the development of a synchronous meiosis induction method with high spore viability. Moreover, our system allows for the shutdown of gene expression and the generation of conditional loss-of-function mutants. This system provides a versatile and powerful tool for manipulating gene expression in fission yeast.

**Summary statement:** A new inducible expression system for fission yeast enhances control, offers compatibility with non-dividing cells, and enables synchronous meiosis induction and conditional loss-of-function analysis.

## Introduction

Understanding gene function is a central pursuit in biological studies. One important approach to dissecting the function of individual genes is through the artificial manipulation of gene expression levels. Over the past 35 years, a large number of regulatable promoter systems have been developed to enable the manipulation of gene expression in the fission yeast *Schizosaccharomyces pombe*. These systems include glucose-repressible promoters (Hoffman and Winston, 1989; Iacovoni et al., 1999), hormone-inducible promoters (Ohira et al., 2017; Picard et al., 1990), thiamine-repressible promoters and their derivatives (Basi et al., 1993; Garg, 2020; Kjaerulff and Nielsen, 2015; Kumar and Singh, 2006; Maundrell, 1993, 1990; Moreno et al., 2000), tetracycline-inducible promoters (Erler et al., 2006; Faryar and Gatz, 1992; Patterson et al., 2019; Zilio et al., 2012), a copper-repressible promoter (Bellemare et al., 2001), a heat shock-inducible promoter (Fujita et al., 2006), a uracil-inducible promoter (Watson et al., 2011; Watt et al., 2008), an ethanol-inducible promoter (Matsuzawa et al., 2013), and pheromone-inducible promoters (Hennig et al., 2018).

Among the various regulatable promoters available for *S. pombe*, the most popular ones are three thiamine-repressible promoters based on that of the *nmt1* gene (Basi et al., 1993; Maundrell, 1993, 1990). This promoter system offers several advantages. Firstly, the *nmt1* promoter (*Pnmt1*) is a highly active promoter when induced, as *nmt1* ranks as the eighth most highly transcribed protein-coding gene in vegetative cells (340 mRNA molecules per cell) (Marguerat et al., 2012). Secondly, there are two weakened promoter variants, *P41nmt1* and *P81nmt1*, which allow for lower levels of expression, approximately one and two orders of magnitude lower than *Pnmt1*, respectively (Basi et al., 1993). Thirdly, these promoters are strongly repressed in the presence of thiamine, resulting in an around 100-fold difference in expression level between the induced and repressed states (Basi et al., 1993; Vještica et al., 2020). However, this system is not without limitations. It is not compatible with yeast extract-based rich media that contain thiamine. Additionally, induction requires a medium switch, which incurs time and labor costs. The most significant drawback is that the intracellular thiamine concentration needs to be diluted through several cell doublings before induced expression can occur (Maundrell, 1990; Tommasino and Maundrell, 1991). This results in a long lag time of around 10-12 h and makes the system unsuitable for non-dividing cells such as cells in a quiescent state.

Tetracycline-inducible systems based on the Tet repressor-operator system of the bacterial transposon Tn*10* have been widely used to manipulate gene expression in eukaryotes (Berens and Hillen, 2003; Hillen et al., 1984, 1983). A basic version of the tetracycline-inducible systems consists of three components: the *tet* operator DNA sequence (*tetO*), which is incorporated into a promoter; the Tet repressor protein (TetR), which has a high binding affinity to *tetO*; and a small-molecule inducer that prevents TetR from binding to *tetO*. Commonly used inducers include tetracycline and its analogs anhydrotetracycline (ahTet) and doxycycline. In the absence of the inducer, TetR binds to *tetO* and represses the transcription of the downstream gene. When the inducer is added, TetR no longer binds to *tetO*, and the transcription of the downstream gene is induced.

Three different tetracycline-inducible promoters have been used in fission yeast. In 1992, Faryar and Gatz established the first tetracycline-inducible promoter for fission yeast by incorporating three *tetO* sequences into the cauliflower mosaic virus 35S (CaMV35S) promoter (Faryar and Gatz, 1992). This tetracycline-inducible promoter can be quickly induced, with a reporter protein product becoming detectable within two hours after the addition of an inducer (Faryar and Gatz, 1992). However, an independent study showed that this system has an induction ratio of only 10-fold and the induced expression level is low, similar to that of *P81nmt1* (Forsburg, 1993). In 2012, Zilio et al. adapted a tetracycline-inducible system, originally developed for the budding yeast *Saccharomyces cerevisiae*, for use in the fission yeast (Bellí et al., 1998; Garí et al., 1997; Zilio et al., 2012). The promoter in this system is composed of seven *tetO* sequences and the basal promoter of the budding yeast *CYC1* gene. The induced level of expression of this promoter is also low, similar to that of *P81nmt1* (Zilio et al., 2012). In 2019, a much stronger tetracycline-inducible promoter, called the *enoTet* promoter (*PenoTet*), was developed by placing three *tetO* sequences into the promoter of the *S. pombe eno101* gene (Patterson et al., 2019). The *eno101* gene is the fourth most highly transcribed protein-coding gene in vegetative cells (470 mRNA molecules per cell) (Marguerat et al., 2012). *PenoTet* achieves an induced expression level about 50% as high as that of *Pnmt1* (Patterson et al., 2019). Unlike the *nmt1* series promoters, these tetracycline-inducible promoters are compatible with rich media, do not need a medium switch for induction, do not exhibit an obvious lag time, and can be used in non-dividing cells. However, these promoters have not been widely adopted, potentially due to the lack of multi-level options, insufficient induced expression levels, non-ideal induction ratios, and the inconvenience associated with the need to introduce both a tetracycline-inducible promoter and a TetR-expressing cassette into a host strain.

In the present study, we have developed an improved tetracycline-inducible expression system for *S. pombe*. This system includes a strong promoter, *PenotetS*, which, similar to *PenoTet*, is derived from the *eno101* promoter but contains only a single *tetO* sequence. *PenotetS* has a high induction ratio of approximately 200-fold and its induced level is similar to that of *Pnmt1*. Additionally, our system includes four weakened variants of *PenotetS*, allowing for five different levels of induced expression. For ease of use, a collection of integration plasmids was constructed, each containing an *enotetS* series promoter and a TetR-expressing cassette. Capitalizing on the compatibility of this system with non-dividing cells, we have developed a new method to induce meiosis, resulting in favorable spore viability and synchrony. Moreover, we have successfully employed this system to generate conditional lethal mutants of essential genes, underscoring its versatility and applicability.

## Results

### The development of a strong inducible promoter *PenotetS*

It has been shown that in the case of the *GAL1* promoter of *Saccharomyces cerevisiae*, placing a single *tetO* sequence between the TATA box and the transcription start site (TSS) is sufficient to render the promoter strongly repressible by TetR (Murphy et al., 2007). We adopted this approach to modify the 276-bp promoter of the *S. pombe eno101* gene (Wang et al., 2014), which was the basis of the *PenoTet* promoter that contains three *tetO* sequences (Patterson et al., 2019). In *S. pombe* promoters, the TATA box and the TSS, each possessing distinct sequence characteristics, are typically separated by a constant distance (Li et al., 2015; Lu and Lin, 2021; Russell, 1983; Thodberg et al., 2019). To maintain this distance, we substituted a 19-bp sequence between the TATA box and the TSS of the *eno101* promoter with a 19-bp *tetO* sequence (Fig. 1A). We named the modified promoter *PenotetS*, with the letter “*S*” indicating the presence of a single copy of the *tetO* sequence.

**Fig. 1.**
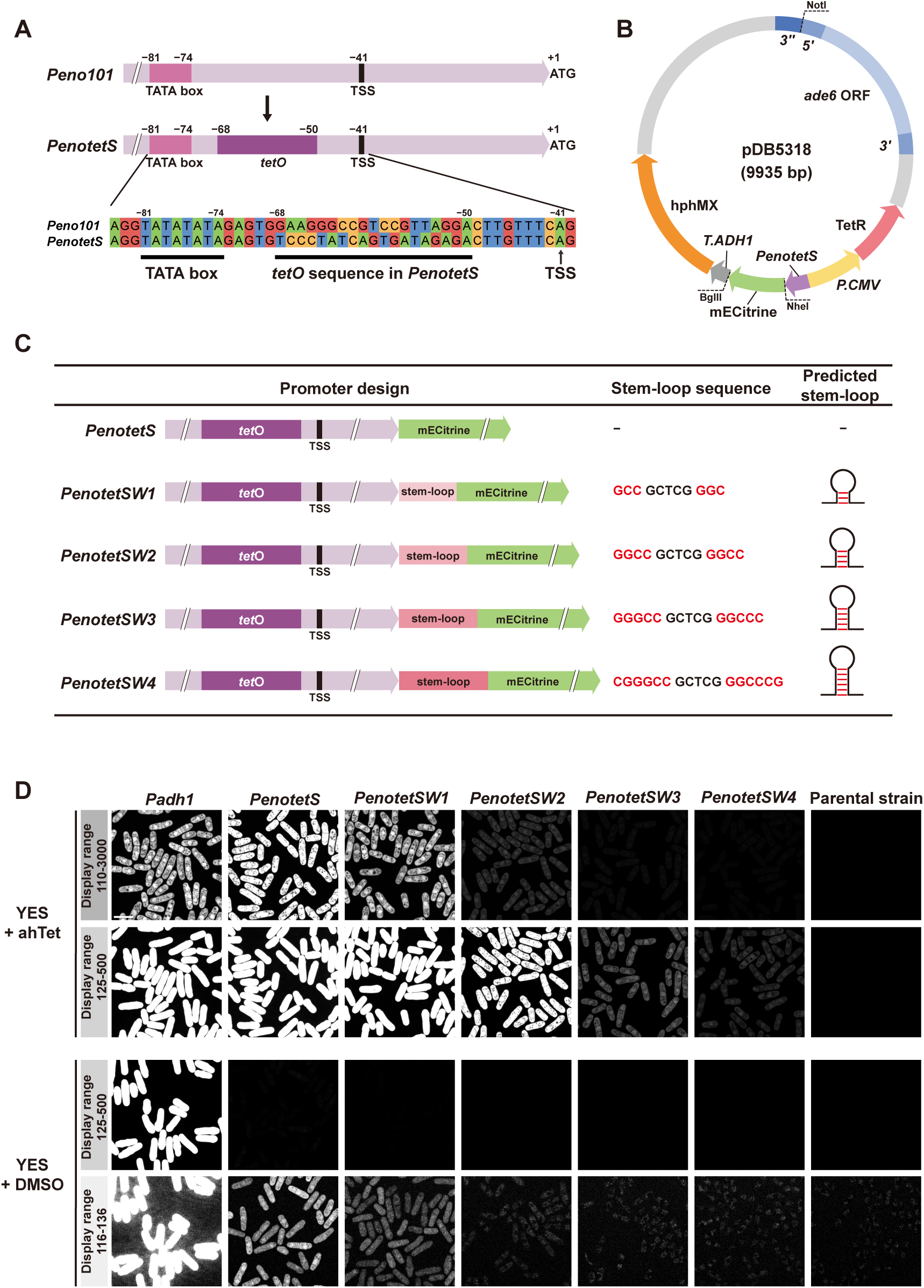
Overview of the new tetracycline-inducible expression system. (A) Schematic showing how the *enotetS* promoter (*PenotetS*) was generated. *PenotetS* was engineered from the *eno101* promoter (*Peno101*) by introducing a single 19-bp *tetO* sequence between the TATA box and the transcription start site (TSS) of *Peno101*, while maintaining the original distance between the TATA box and the TSS. (B) Diagram of the integration plasmid pDB5318. This plasmid contains *PenotetS* and a TetR-expressing cassette, which consists of the cytomegalovirus promoter (*P.CMV*) and the TetR coding sequence. The plasmid can be integrated at the *ade6* locus following linearization with NotI digestion. The mECitrine coding sequence can be replaced with the sequence of any desired gene to be expressed under the control of *PenotetS*, using the NheI and BglII restriction sites. (C) Schematic of four weakened variants of *PenotetS*. These variants were created by inserting a GC-rich stem-loop-forming sequence downstream of *PenotetS*. Bases forming the stem are highlighted in red, while those forming the loop are shown in black. (D) Fluorescence of cells expressing the yellow fluorescent protein mECitrine from each of the *enotetS* series promoters and from the *adh1* promoter in the presence and absence of the inducer anhydrotetracycline (ahTet). Cells were imaged after 24 hours of growth in YES medium in the presence of ahTet or DMSO (solvent control). The images are presented with two different display ranges to enhance the visualization of differences in fluorescence intensities. Scale bar, 10 μm.

A drawback of some of the previously published tetracycline-inducible systems for *S. pombe* is the inconvenience caused by the need to separately introduce a tetracycline-inducible promoter and a TetR-expressing cassette. To establish a user-friendly system, we combined *PenotetS* with a TetR-expressing cassette, which consists of the cytomegalovirus (CMV) promoter and the TetR coding sequence, in a stable integration vector (SIV) (Vještica et al., 2020) (Fig. 1B). The CMV promoter is a robust constitutive promoter in *S. pombe* (Amelina et al., 2016; Toyama and Okayama, 1990). Additionally, downstream of *PenotetS*, we inserted the coding sequence for the yellow fluorescent protein mECitrine, followed by the terminator sequence derived from the *ADH1* gene of *Saccharomyces cerevisiae*. The resulting plasmid was named pDB5318. The fluorescence emitted by mECitrine serves as an indicator of the expression level of *PenotetS*.

In *S. pombe*, previous studies have shown that anhydrotetracycline (ahTet), an analog of tetracycline, is a more effective inducer compared to tetracycline and doxycycline (Erler et al., 2006; Zilio et al., 2012). When an *S. pombe* strain containing pDB5318 was cultured in YES medium supplemented with 2.5 μg/mL of ahTet, a concentration used in a previous study (Zilio et al., 2012), it was observed that the expression level of *PenotetS* was higher than that of the strong constitutive promoter *Padh1* (Fig. 1D). In the absence of ahTet, the expression level of *PenotetS* was markedly lower than that of *Padh1*. Thus, *PenotetS* is an inducible promoter with a high level of induced expression and a substantial induction ratio. To determine whether the expression of TetR or treatment with ahTet affects the growth rate, we conducted a growth curve analysis using a strain containing pDB5318 and a control strain lacking pDB5318 (Fig. S1). The growth rates of both strains were identical in media with or without ahTet, indicating that neither the expression of TetR nor the presence of 2.5 μg/mL of ahTet in the culture medium impacted the cell growth rate.

### The development of four weakened variants of *PenotetS*

To provide a range of different expression levels, we developed four weakened variants of *PenotetS*. The design was inspired by a previous study that used GC-rich stem-loop-forming sequences inserted in the 5′ UTR to reduce gene expression (Lamping et al., 2013). It was shown that the extent of expression attenuation increases with the number of GC base pairs in the stem. We selected four different stem-loop-forming sequences used in that study (Fig. 1C). Each sequence was inserted immediately downstream of *PenotetS*. The resulting promoters were named *PenotetSW1*, *PenotetSW2*, *PenotetSW3*, and *PenotetSW4*. The letter “*W*” indicates a weakened variant, while the number denotes the expected expression strength, with *PenotetSW1* being the highest and *PenotetSW4* being the lowest.

These weakened promoters were combined with the TetR-expressing cassette in an SIV plasmid, following the same procedure as *PenotetS*. In YES medium supplemented with 2.5 μg/mL of ahTet, the expression strengths of these promoters varied as expected (Fig. 1D). *PenotetSW1* exhibited an expression level similar to that of *Padh1*, while the other three promoters showed progressively lower levels. In the absence of ahTet, the expression level of each promoter was dramatically lower compared to when ahTet was present, with the three weaker promoters showing fluorescence signals that were indistinguishable from the background.

### Inducer dose response

We assessed the inducer dose response of *PenotetS* in both YES and EMM media. The inducer ahTet was used at a range of six concentrations (0, 0.004, 0.02, 0.1, 0.5, and 2.5 μg/mL). After a 24-hour induction period (which was long enough to reach a steady-state level; see below), the expression levels were measured using the fluorescence signal of the reporter protein mECitrine (Fig. 2). In both types of media, a dramatic transition from a near-minimal level to a near-maximal level was observed between two doses that differed by 5-fold. In YES, this transition occurred between 0.004 μg/mL and 0.02 μg/mL of ahTet (Fig. 2B), while in EMM, it occurred between 0.02 μg/mL and 0.1 μg/mL of ahTet (Fig. 2D). Similar steep dose-response patterns have been observed in other published eukaryotic tetracycline-inducible systems that utilize constitutively expressed TetR as a repressor (Azizoglu et al., 2021; Erler et al., 2006; Nevozhay et al., 2009). We chose to use 2.5 μg/mL of ahTet, an inducer concentration well above the saturating level in both types of media, for subsequent experiments.

**Fig. 2.**
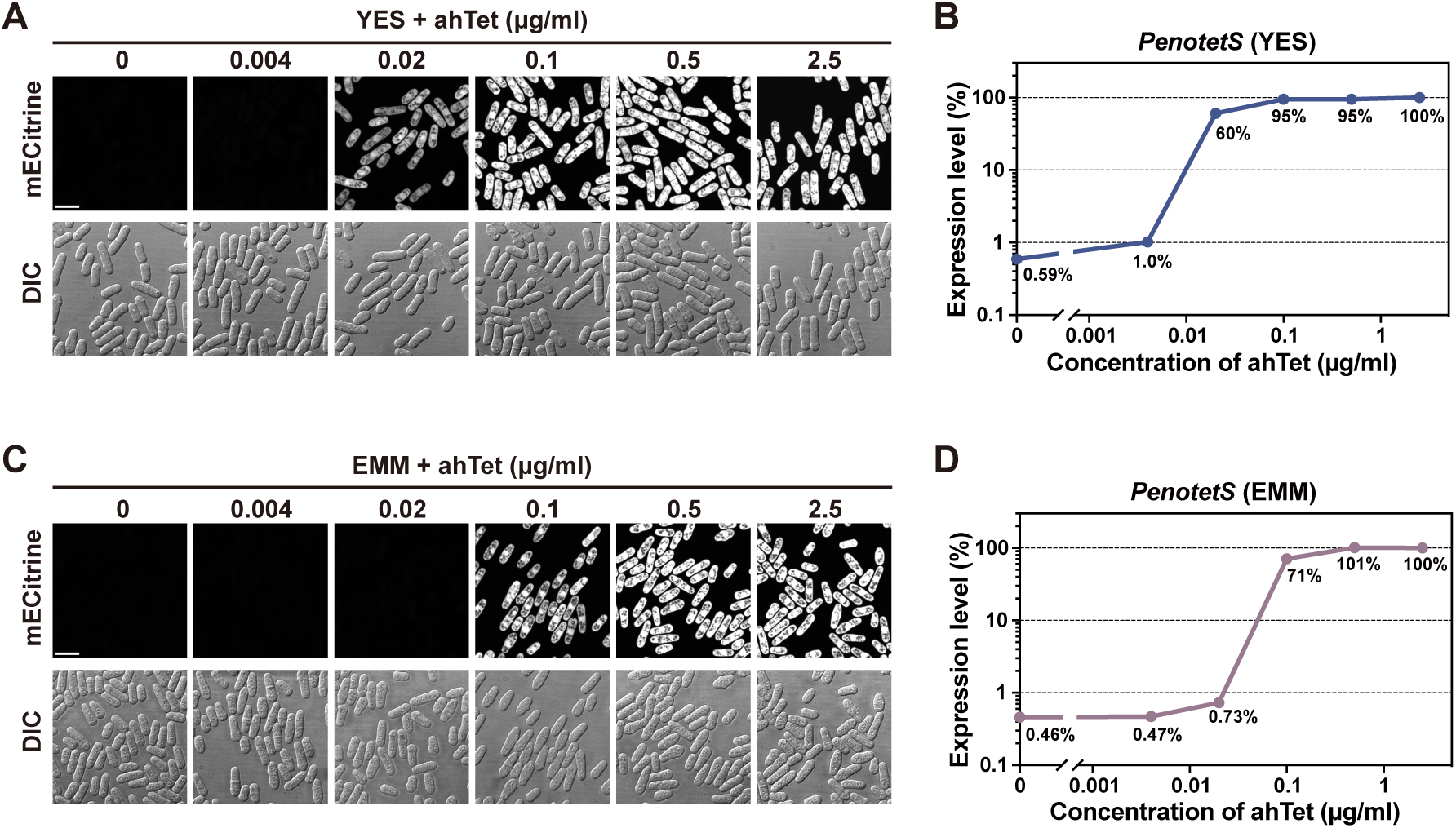
Inducer dose response of *PenotetS*. (A) Inducer dose response in YES medium. Cells expressing mECitrine from *PenotetS* were grown in YES medium in the presence of different doses of the inducer ahTet and then imaged. Scale bar, 10 μm. (B) Quantification of mECitrine fluorescence in the imaging data from the experiment shown in A. The fluorescence level in the presence of 2.5 μg/ml ahTet was used as a reference for normalization (100% expression level). (C) Inducer dose response in EMM medium. Cells expressing mECitrine from *PenotetS* were grown in EMM medium in the presence of different doses of the inducer ahTet and then imaged. Scale bar, 10 μm. (D) Quantification of mECitrine fluorescence in the imaging data from the experiment shown in C. The fluorescence level in the presence of 2.5 μg/ml ahTet was used as a reference for normalization (100% expression level).

### Induction kinetics in dividing cells

We next assessed the induction kinetics of *PenotetS* and *PenotetSW2* by performing induction time course experiments in both YES and EMM media (Fig. 3). In all time courses, the expression level of the reporter protein mECitrine increased by more than two orders of magnitude within the first 10 hours following the addition of ahTet and ceased to increase after 12 hours. We used the expression level at the 24-hour time point as the steady-state level. In YES medium, for both promoters, at the one-hour time point after adding ahTet to the cultures, the fluorescence signal of mECitrine had already increased by more than one order of magnitude, reaching around 10% of the steady-state level. This rapid increase suggests that both promoters were turned on quickly after the addition of ahTet. In EMM medium, the initial phase of the induction was slightly slower, reaching around 5% of the steady-state level at the one-hour time point. In both types of media, at the four-hour time point, both promoters had reached or surpassed 50% of the steady-state level. By the eight-hour time point, both promoters had reached within 15% of the steady-state level.

**Fig. 3.**
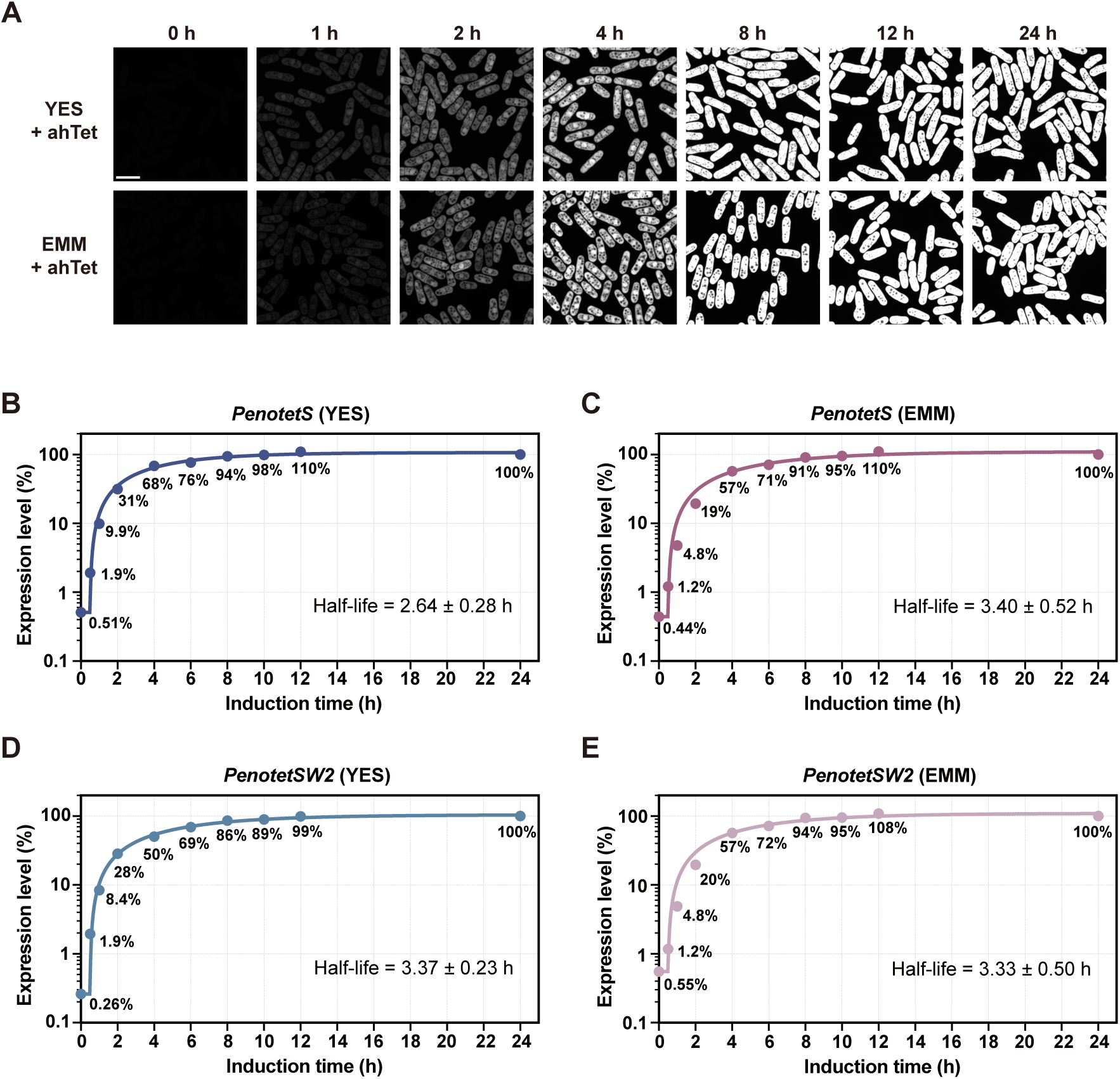
Induction kinetics of *PenotetS* and *PenotetSW2* in dividing cells. (A) Time-course analysis of the induction kinetics of *PenotetS* in log-phase cells grown in YES and EMM media. Images of cells expressing mECitrine from *PenotetS* were taken at the indicated time points after the addition of 2.5 μg/ml ahTet to the culture. Scale bar, 10 μm. (B, C) Quantification of mECitrine fluorescence in the imaging data from the experiment shown in A. The fluorescence level at the 24-hour time point was used as a reference for normalization (100% expression level). The curves depict the mathematical model of the induction kinetics, with the modeled half-life value presented beneath each curve. (D, E) Induction kinetics of *PenotetSW2* in log-phase cells grown in YES and EMM media. Data processing and presentation were carried out as described in (B, C).

Upon the activation of an inducible promoter that drives the expression of a protein-coding gene, the level of the protein product reaches a steady state when its synthesis is balanced by its degradation and dilution (Azizoglu et al., 2021; Rosenfeld et al., 2002). The steady-state expression level is determined by both the synthesis rate and the degradation-and-dilution rate, while the induction kinetics are solely determined by the degradation-and-dilution rate (Rosenfeld et al., 2002). Fluorescent proteins are highly stable and have in vivo half-lives longer than 20 hours (Corish and Tyler-Smith, 1999; He et al., 2019; Perruca-Foncillas et al., 2022). In our experimental setup, given that the doubling time of proliferating *S. pombe* cells is around 3 hours (Petersen and Russell, 2016), the degradation-and-dilution rate of mECitrine should mainly be determined by its dilution through cell division. In this scenario, without considering any delay between transcription activation and the formation of mECitrine protein capable of emitting fluorescence, the time required to reach 50% of the steady-state expression level theoretically equals the doubling time (Rosenfeld et al., 2002). However, nascent fluorescent proteins only become fluorescent after undergoing a maturation process, which takes approximately 30 minutes for mECitrine (Azizoglu et al., 2021). Taking this delay into account, the induction kinetics observed in our experiments align closely with immediate or near-immediate transcription activation upon the addition of ahTet. To better characterize the induction kinetics, we performed quantitative modeling by fitting the experimental data to an induction-kinetics model. The resulting fitted curves closely align with our quantitative data, and the half-life parameters derived from the modeling are consistent with our proposed explanation of the expression kinetics (Fig. 3B-D). This analysis further supports the conclusion that transcription activation occurs rapidly upon the addition of ahTet.

### Quantification of expression levels and comparison with the *nmt1* series promoters

Next, we conducted a comprehensive quantitative analysis of the induced and repressed expression levels of all *enotetS* series promoters in YES and EMM media, and made comparisons with *Padh1* in YES medium and the *nmt1* series promoters in EMM medium (Fig. 4). In YES medium, the induced expression level of *PenotetS* was approximately 1.6-fold that of *Padh1* (Fig. 4A). The fold differences between the induced level of *PenotetS* and the induced levels of the four weakened *enotetS* series promoters were approximately 1.7-fold, 7.1-fold, 30-fold, and 54-fold, respectively (Fig. 4A). The fold differences between the induced and repressed levels (induction ratios) were about 207-fold for *PenotetS* and about 226-fold for *PenotetSW1* (Fig. 4A). Since the repressed levels of *PenotetSW2*, *PenotetSW3*, and *PenotetSW4* were very close to the background, we could only determine that these levels were below a threshold (THR) for accurate measurement (Fig. 4A).

**Fig. 4.**
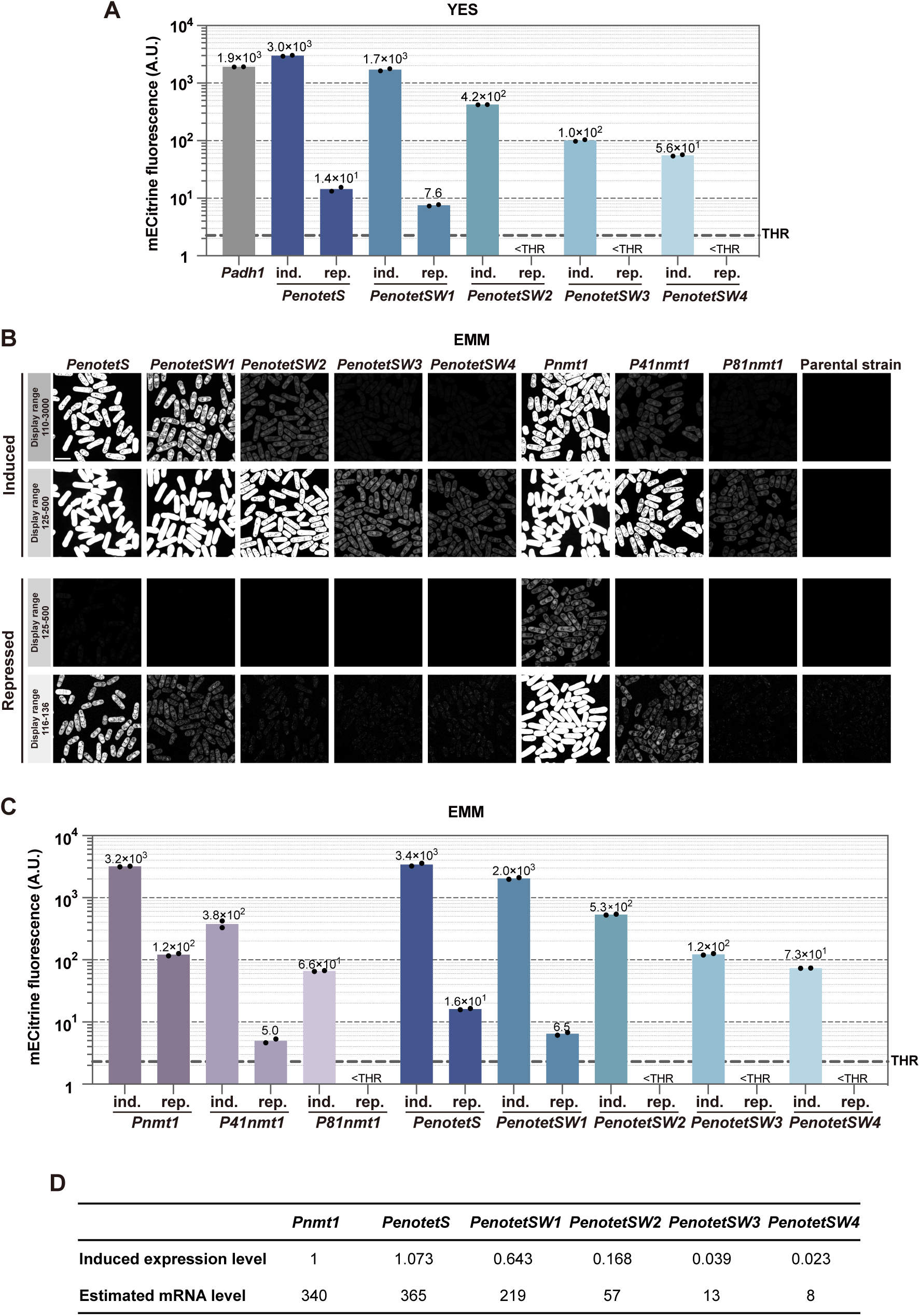
Expression levels of different promoters. (A) Expression levels in YES medium. Cells expressing mECitrine from *Padh1* and from each of the *enotetS* series promoters were grown in YES medium under induced conditions (abbreviated as ind., in the presence of ahTet) and repressed conditions (abbreviated as rep., in the absence of ahTet), and then imaged. The quantification results from two biological replicates and their mean values are shown. A thick dashed line represents a threshold (THR) below which the mECitrine fluorescence level cannot be accurately determined. This threshold corresponds to 2% of the average background fluorescence in a parental strain lacking mECitrine. (B) Expression levels in EMM medium. Cells expressing mECitrine from each of the *enotetS* series promoters were grown in EMM medium under induced conditions (in the presence of ahTet) and repressed conditions (in the absence of ahTet), and then imaged. Cells expressing mECitrine from each of the *nmt1* series promoters were grown in EMM medium under induced conditions (in the absence of thiamine) and repressed conditions (in the presence of thiamine), and then imaged. The images are presented with two different display ranges to enhance the visualization of differences in fluorescence intensities. Scale bar, 10 μm. (C) Quantification of mECitrine fluorescence in the imaging data from the experiment shown in B. The quantification results from two biological replicates and their mean values are shown. (D) The induced expression level based on the mECitrine fluorescence (normalized to that of *Pnmt1*) and the estimated number of mRNA molecules per cell based on the reported value of 340 for the number of mRNA molecules per cell for the *nmt1* gene.

In EMM medium, the induced and repressed expression levels of the *enotetS* series promoters mirrored those observed in YES medium (Fig. 4B,C). When compared to the *nmt1* series promoters, the induced expression level of *PenotetS* was almost the same as *Pnmt1* and the induced levels of *PenotetSW2* and *PenotetSW4* were similar to those of *Pnmt41* and *P81nmt1*, respectively (Fig. 4B,C). Therefore, in comparison to the *nmt1* series promoters, the *enotetS* series promoters span the same range of induced levels but provide more intermediate levels. Using the mRNA level of the *nmt1* gene in vegetative cells (340 mRNA molecules per cell) as a reference (Marguerat et al., 2012; Rutherford et al., 2024), the induced expression levels of *PenotetS*, *PenotetSW1*, *PenotetSW2*, *PenotetSW3*, and *PenotetSW4* correspond to approximately 365, 219, 57, 13, and 8 mRNA molecules per cell, respectively (Fig. 4D). It is important to note that mRNA levels do not necessarily correlate with protein levels, and the expression of different proteins from the same promoter can result in markedly different protein levels. Regarding repressed expression levels, the *enotetS* series promoters had lower repressed levels than the *nmt1* series promoters when comparing promoters with similar induced levels. In our experiments, *Pnmt1* exhibited only a 26-fold induction ratio, which is lower than the reported induction ratio of around 100-fold (Basi et al., 1993; Vještica et al., 2020). The reason behind this discrepancy is unclear. Nevertheless, even when considering the reported induction ratio for comparison, the *enotetS* series promoters still outperformed the *nmt1* series promoters in this aspect.

### Induction kinetics and expression levels in quiescent cells

One limitation of the *nmt1* series promoters is their unsuitability for non-dividing cells, as the induction of these promoters requires the dilution of intracellular thiamine through multiple rounds of cell division after shifting the cells to a thiamine-free medium. Tetracycline-inducible systems should be free of this limitation. Therefore, we tested whether *PenotetS* can function in two types of non-dividing cells: cells arrested in the G0 cell cycle phase through nitrogen starvation and spores (Fig. 5).

**Fig. 5.**
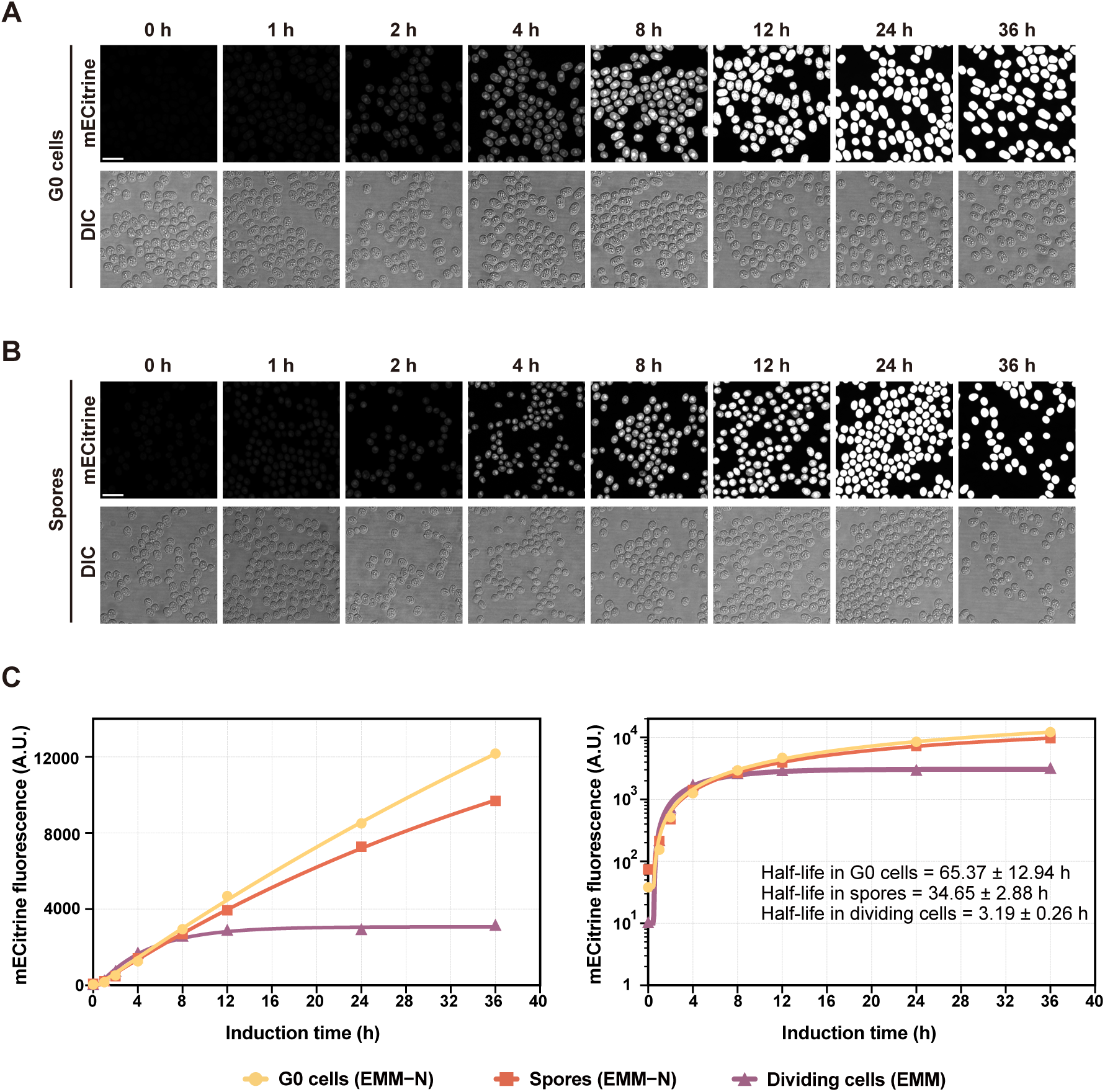
Induction kinetics of *PenotetS* in quiescent cells. (A) Time-course analysis of the induction kinetics of *PenotetS* in G0 cells in EMM−N medium. Images were taken at the indicated time points after the addition of 2.5 μg/ml ahTet. Scale bar, 10 μm. (B) Time-course analysis of the induction kinetics of *PenotetS* in spores in EMM−N medium. Images were taken at the indicated time points after the addition of 2.5 μg/ml ahTet. Scale bar, 10 μm. (C) Quantification of mECitrine fluorescence in the imaging data from the experiments shown in A and B, as well as a control experiment in which log-phase cells were grown in EMM medium. Three biological replicates were conducted, and one representative dataset is shown here. The graph on the left uses a linear scale for the y-axis, while the graph on the right uses a logarithmic scale. The curves depict the mathematical model of the induction kinetics, with the modeled half-life values presented in the graph on the right.

The basal expression levels of *PenotetS* in G0 cells and spores, as reported by the fluorescence of mECitrine, were slightly higher than the basal level in the dividing cells (Fig. 5C), presumably due to the lack of dilution of mECitrine through cell division. Upon the addition of ahTet to G0 cells and spores, the expression of mECitrine from *PenotetS* increased several-fold within the first hour and reached a level similar to the steady-state level in the dividing cells at the eight-hour time point. Unlike the situation in the dividing cells, where the expression of mECitrine from *PenotetS* reached a steady state at the 12-hour time point, the level of mECitrine continued to increase in G0 cells and spores after the 12-hour time point (Fig. 5C). The increase did not stop after the 24-hour time point and the level of mECitrine reached several-fold higher than the steady-state level in the dividing cells at the 36-hour time point (Fig. 5C). The prolonged rise time and the higher induced levels in G0 cells and spores can be explained by the degradation-and-dilution rate of mECitrine no longer being determined by its dilution through cell division but rather by the degradation of mECitrine, which has a long half-life.

### Meiosis induction in *h+/h+* diploids using *PenotetSW1*-expressed Mc-2A-Mi

In *S. pombe,* the most commonly used method to induce synchronous meiosis is through the inactivation of a negative regulator of meiosis, Pat1, using a temperature-sensitive allele of the *pat1* gene, *pat1-114*, in diploid cells homozygous for the mating type (*h+/h+* or *h−/h−*) (Bähler et al., 1991; Iino and Yamamoto, 1985). However, this approach has several drawbacks, including the use of non-physiological temperature and low spore viability (Bähler et al., 1991; Chikashige et al., 2004; Cipak et al., 2012; Guerra-Moreno et al., 2012; Yamamoto and Hiraoka, 2003). The development of ATP analog-sensitive alleles of *pat1* has enabled the induction of synchronous meiosis at physiological temperature (Cipak et al., 2012; Guerra-Moreno et al., 2012). Moreover, combining Pat1 inactivation with the expression of the *matP*-cassette-encoded protein Pc in *h−/h−* diploids has been shown to elevate spore viability to near-wild-type levels (Cipak et al., 2012; Yamamoto and Hiraoka, 2003). Thus, the current state-of-the-art approach for achieving synchronous meiosis with normal spore viability in *S. pombe* requires the combination of two genetic modifications. To showcase the practical utility of our *enotetS* series promoters, we devised a single-plasmid-based method for inducing synchronous meiosis.

Diploid *S. pombe* cells that are heterozygous for the mating type (*h+/h−*) can undergo meiosis, whereas those that are homozygous for the mating type (*h+/h+* or *h−/h−*) cannot. This is because the latter lack proteins encoded by one of the two types of mating-type cassettes. In the case of *h+/h+* diploid cells, the missing proteins are the *matM*-cassette-encoded proteins Mc and Mi (Kelly et al., 1988). In our new method for inducing meiosis, we used *PenotetSW1* to induce the expression of Mc and Mi linked by a self-processing 2A peptide (Mc-2A-Mi) in *h+/h+* diploid cells (Fig. 6A). *h+/h+* cells harboring an integration plasmid expressing Mc-2A-Mi under the control of *PenotetSW1* were initially cultured in YES medium and subsequently transferred to EMM−N medium to arrest the cells in G0. Following a 24-hour incubation in EMM−N medium, ahTet was added to induce meiosis. Time course analysis demonstrated the successful induction of synchronous meiosis (Fig. 6B). Tetrad analysis revealed a spore viability of 97% for meiosis induced using this method (Fig. 6C). Thus, we have established a new method for inducing meiosis that is characterized by good synchrony and normal spore viability.

**Fig. 6.**
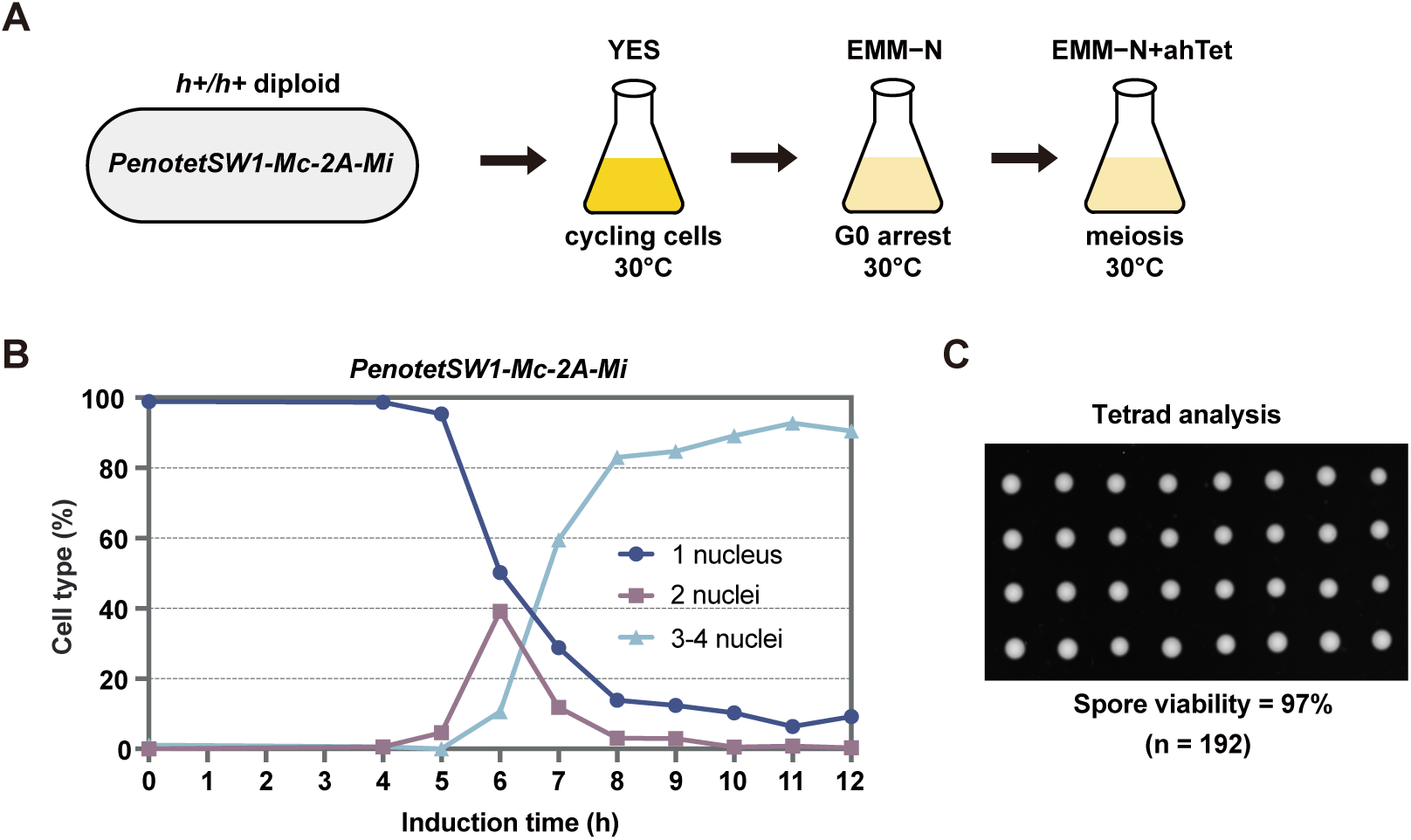
A new method for synchronous meiosis induction based on the expression of *Mc-2A-Mi* from *PenotetSW1* in *h+/h+* diploids. (A) Schematic illustrating the new method for synchronous meiosis induction. (B) Progression of meiosis monitored by DNA staining. Cell samples were stained with the DNA dye SYTOX Green and imaged at the indicated time points after the addition of ahTet. Manual counting was performed to determine the percentages of cells containing different numbers of nuclei. Three biological replicates were conducted, and one representative dataset is shown here. (C) Viability of spores from induced meiosis. Tetrad analysis was performed on YES plates.

### Using *enotetS* series promoters to create conditional loss-of-function mutants

Studying the functions of essential genes relies on the creation of conditional loss-of-function mutants. Previously, we developed an auxin-inducible degron (AID) system for generating such mutants in *S. pombe* (Zhang et al., 2022). However, not all genes are compatible with the AID approach. In an unrelated project, we applied the AID method to construct conditional loss-of-function mutants of three essential genes: *ypp1*, *sui1*, and *nrs1*. We first performed endogenous AID tagging in a diploid background and obtained heterozygous tagged diploid strains. Then, we attempted to derive haploid AID-tagged progeny through tetrad dissection (Fig. S2). AID-tagged haploid progeny were readily obtained for *ypp1*. However, no viable AID-tagged haploid progeny were obtained for *sui1* and *nrs1*, possibly due to AID tagging interfering with the function of the protein products of these two genes. We decided to take this opportunity to explore the feasibility of using the *enotetS* series promoters for generating conditional loss-of-function mutants, especially for genes that are not amenable to the AID approach.

We first separately deleted *sui1* and *nrs1* in a diploid background to obtain heterozygous deletion diploid strains. Then, for each gene, a plasmid expressing the gene from *PenotetSW2* and a plasmid expressing the gene from *PenotetSW3* were separately integrated into the heterozygous deletion diploid strain. These two promoters were chosen because their induced expression levels are close to the mRNA levels of *sui1* and *nrs1* in vegetative cells (11 and 20 mRNA molecules per cell, respectively) (Marguerat et al., 2012; Rutherford et al., 2024). The resulting strains were subjected to tetrad analysis on YES plates with and without ahTet (Fig. 7). On YES plates without ahTet, for both genes, the basal level expression from neither promoter was able to allow the deletion haploids to form colonies (Fig. 7A). In contrast, on YES plates with ahTet, for both genes, the induced expression from either promoter rescued the inviability of the deletion haploids (Fig. 7A). *PenotetSW2*-driven induced expression in deletion haploids resulted in colony growth indistinguishable from wild type, whereas *PenotetSW3*-driven induced expression resulted in colonies slightly smaller than wild type at 3 days but not at 6 days (Fig. 7A). Thus, we successfully obtained conditional lethal mutants for both genes.

**Fig. 7.**
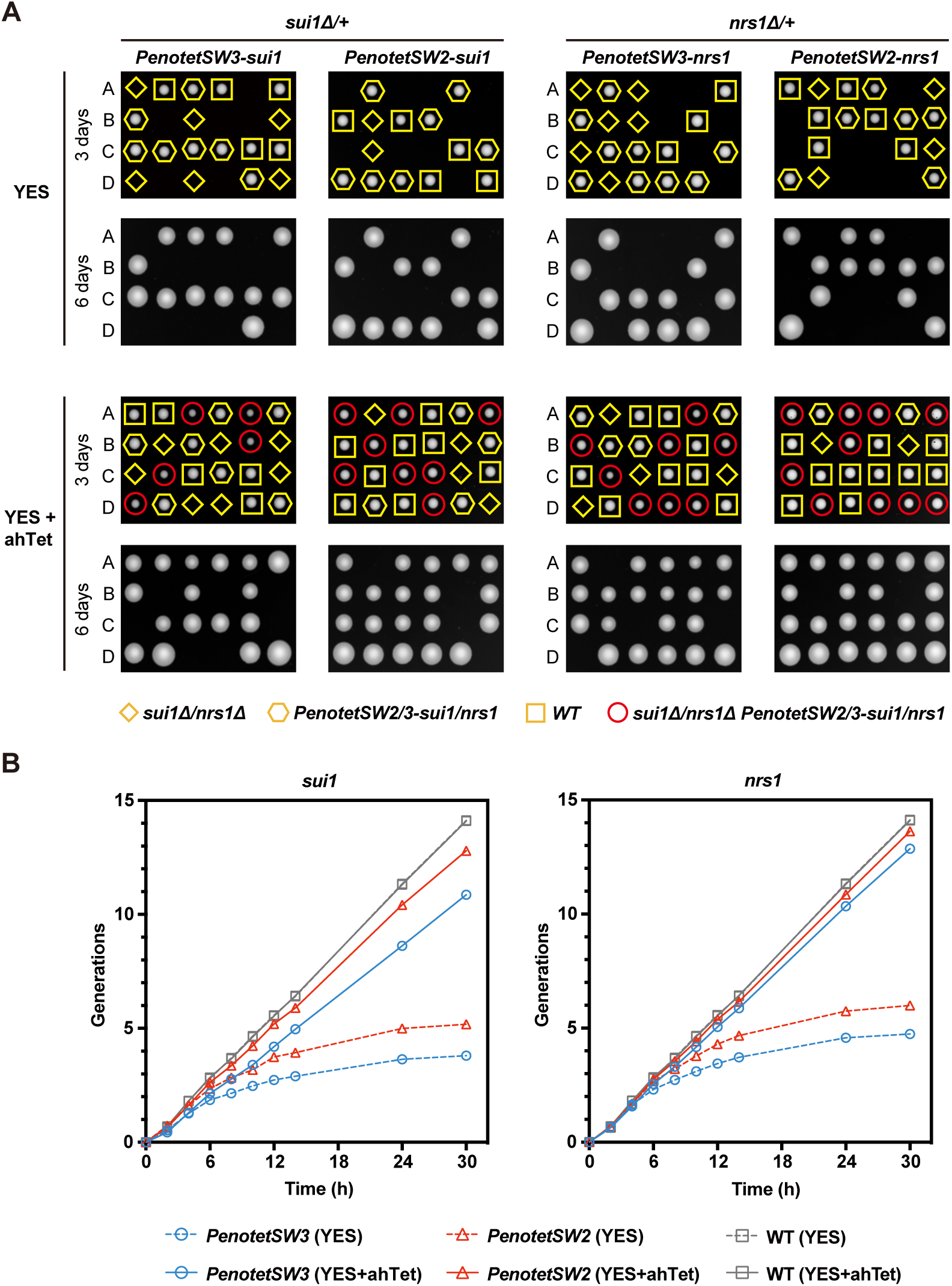
Generation of conditional loss-of-function mutants using the *enotetS* series promoters. (A) Tetrad analysis of heterozygous deletion diploids carrying an integration plasmid expressing the deleted gene from either *PenotetSW2* or *PenotetSW3*. Diploids were sporulated, and the resulting tetrads were dissected on YES plates with and without ahTet. Four spores from a single ascus are labeled A, B, C, and D. The genotype of inviable spores was inferred based on the assumption of 2:2 segregation. Induced expression of *sui1* and *nrs1* rescued the lethality of the respective deletions, while repressed expression did not. (B) Growth curve analysis showing that the growth defect caused by shutting off gene expression through the removal of ahTet arises rapidly. Cells were initially cultured for 24 hours in YES liquid medium containing ahTet. Following this, cells were washed three times with water and transferred to both ahTet-containing and ahTet-free YES media. The optical density at 600 nm (OD600) of each culture was measured at the indicated time points. Cultures were diluted when the OD600 approached 1.0 to maintain logarithmic growth. The number of generations was calculated based on the OD600 measurements and the dilution ratio.

To determine how quickly the loss-of-function phenotype arises upon the removal of ahTet, we conducted a growth curve analysis (Fig. 7B). Haploid deletion strains harboring a plasmid expressing the deleted gene from *PenotetSW2* or *PenotetSW3* were pre-cultured in YES liquid medium with ahTet. After the cells were harvested and washed with water, they were transferred to fresh YES medium with and without ahTet. At the six-hour time point, the cultures in the ahTet-free YES medium started to exhibit a noticeably lower growth rate compared to the cultures in the ahTet-containing YES medium. The differences in growth rate became more pronounced as time elapsed. Given that the loss-of-function phenotype does not manifest immediately after the promoter is turned off, but rather arises only after the protein product has been significantly depleted through dilution and degradation, these findings suggest that the *enotetS* series promoters can be rapidly switched off after shifting cells to an ahTet-free medium. Therefore, these promoters provide useful tools for analyzing loss-of-function phenotypes of essential genes in *S. pombe*.

## Discussion

In this study, we have developed the *enotetS* series promoters, a new set of tetracycline-inducible promoters for *S. pombe* (Fig. 8). These promoters offer a wide range of expression levels and allow for rapid induction of gene expression in both dividing and non-dividing cells. By combining each of these promoters with a TetR-expressing cassette, this system can be easily implemented in *S. pombe* through a single plasmid transformation. To demonstrate the usefulness of this new inducible expression system, we have successfully developed a new method for inducing synchronous meiosis and generated conditional loss-of-function mutants for two essential genes.

**Fig. 8.**
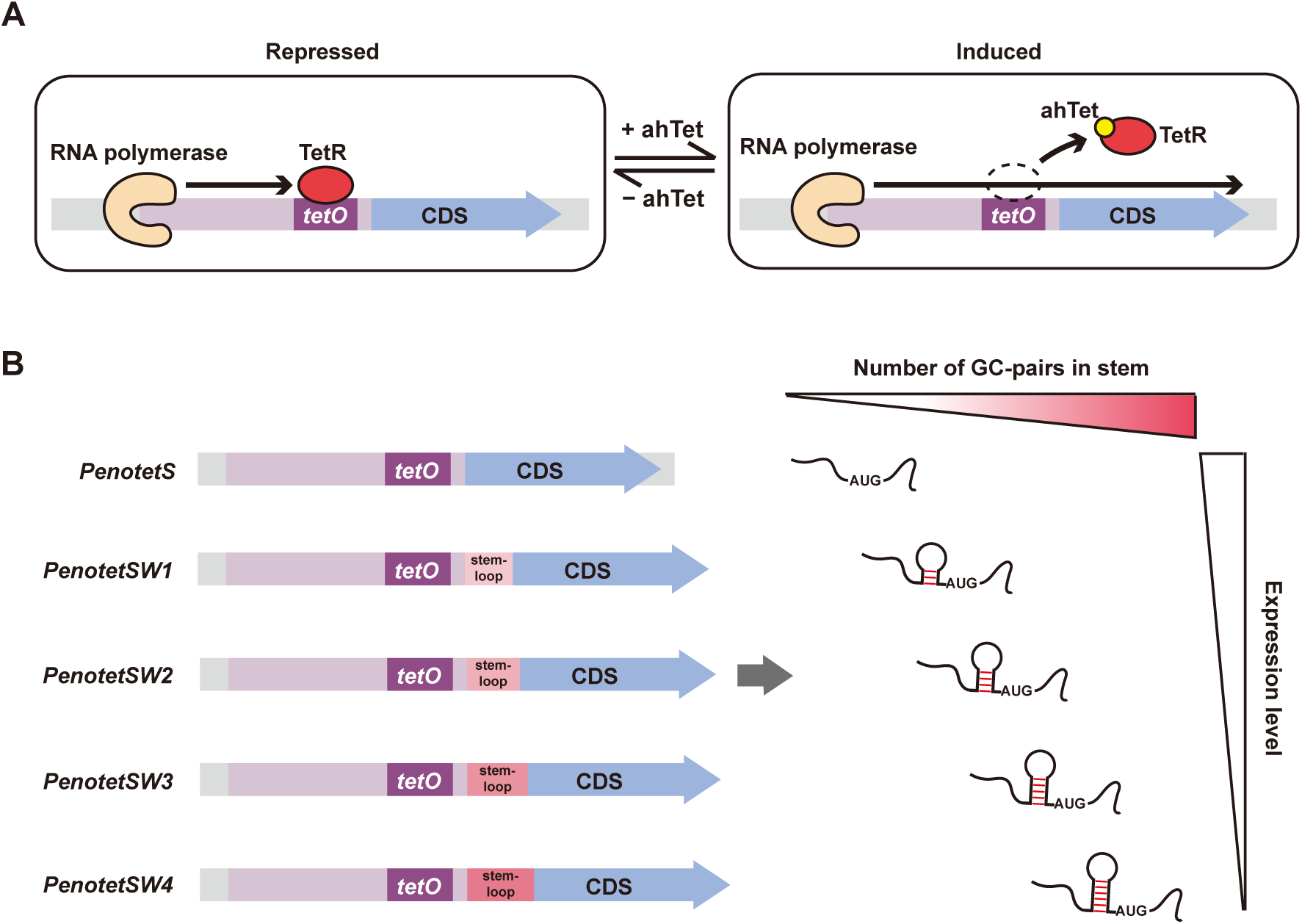
Schematics depicting the improved tetracycline-inducible expression system developed in this study. (A) Schematic illustrating the repressed and induced states of the *enotetS* promoter. In the absence of the inducer ahTet, TetR binds to *tetO*, repressing the transcription of the downstream gene. Upon the addition of ahTet, the ahTet-bound TetR undergoes a conformational change, preventing it from binding to *tetO* and thereby allowing the transcription of the downstream gene to be induced. The induced state can be reverted back to the repressed state by removing ahTet. (B) Schematic illustrating the five distinct expression levels of the *enotetS* series promoters. Four different stem-loop-forming sequences were individually inserted immediately downstream of *PenotetS*, resulting in four weakened variants: *PenotetSW1*, *PenotetSW2*, *PenotetSW3*, and *PenotetSW4*. The degree of expression attenuation increases with the number of GC base pairs in the stem.

Tetracycline and its analogs have traditionally been considered biologically inert to eukaryotic physiology (Gossen et al., 1993). However, recent research has demonstrated that doxycycline, a commonly used inducer in mammalian tetracycline-inducible systems, disturbs mitochondrial protein translation, presumably due to the bacterial origin of mitochondria (Moullan et al., 2015). This raises concerns regarding the use of tetracycline-inducible systems (Chatzispyrou et al., 2015; De Boeck and Verfaillie, 2021; Ohira et al., 2017). Unlike tetracycline and doxycycline, which strongly inhibit bacterial protein translation and belong to the category of “classical tetracyclines”, the inducer used in our system, ahTet, does not effectively inhibit bacterial protein translation and is considered an “atypical tetracycline” (Rasmussen et al., 1991). Therefore, it is unlikely that ahTet negatively affects mitochondrial protein translation. Consistent with this, even though mitochondrial protein translation is essential for *S. pombe* proliferation, studies that have used ahTet on *S. pombe*, including our own, have not observed any detrimental effects on proliferation (Berg and Moseley, 2023; Erler et al., 2006; Lemay et al., 2014; Patterson et al., 2019; Shiroiwa et al., 2011; Sunder et al., 2012; Torres-Garcia et al., 2020; Wang et al., 2018; Zilio et al., 2012). As a precautionary measure against potential unknown side effects of ahTet on *S. pombe*, especially when investigating non-growth phenotypes, we recommend including a strain without an *enotetS* series promoter as a control to rule out the possibility that a phenotype is caused by side effects of ahTet. ahTet is a cost-effective inducer. When used at a concentration of 2.5 μg/mL, the cost of ahTet for making one liter of ahTet-containing medium is about 6 US dollars (based on the list price of Sigma-Aldrich catalog #37919). The ahTet-containing medium should be prepared fresh from the DMSO-dissolved ahTet stock solution before use, as ahTet is not very stable in aqueous solution. It has been reported that ahTet has a half-life of around 30 hours in culture media at 30°C (Politi et al., 2014).

A higher level of ahTet is required for induction in EMM than in YES (Fig. 2), and the induction kinetics appear to be slightly slower at the early stage of induction in EMM than in YES (Fig. 3). We speculate that these differences may be due to a reduced uptake of ahTet in EMM. It has been reported that the addition of 5 mM of Mg^2+^ to the culture medium diminishes the uptake of tetracycline into *E. coli* cells (Zhang et al., 2014). EMM contains a notably higher concentration of Mg^2+^ (5.2 mM) than YES (less than 0.5 mM) (Li et al., 2020; Petersen and Russell, 2016).

The five induced expression levels of the *enotetS* series promoters offer a wide range of possibilities for manipulating gene expression levels. In theory, even more intermediate expression levels can be obtained by titrating the inducer dose. However, the *PenotetS* promoter (and likely its weakened variants) exhibits a steep dose-response curve. This means that the range of inducer doses over which the promoter is responsive, known as the input dynamic range, is narrow. Consequently, precise adjustments to expression levels by altering the inducer dose can be challenging (Azizoglu et al., 2021). If desired, the input dynamic range can be expanded by implementing negative auto-regulation of TetR (Azizoglu et al., 2021; Madar et al., 2011; Nevozhay et al., 2009; Patterson et al., 2019).

The use of the *enotetS* series promoters to generate conditional lethal mutants of essential genes is a valuable alternative to the AID method, especially for genes that cannot tolerate AID tagging. This strategy, known as promoter shut-off, has previously been implemented in *S. pombe* using the *nmt1* series promoters and has successfully revealed the loss-of-function phenotypes of many essential genes (Bestul et al., 2021; Kelly et al., 1993; Matynia et al., 2002; Ottilie et al., 1995; Petersen and Hagan, 2003; Tajadura et al., 2004; Wang et al., 2002; Westwood et al., 2004; Win et al., 2004; Yue et al., 2014). Compared to the *nmt1* series promoters, the *enotetS* series promoters offer more levels of expression and larger ratios between induced and repressed conditions. Therefore, they are expected to be better tools for promoter shut-off.

While we generated the promoter shut-off strains for *sui1* and *nrs1* by first establishing heterozygous deletion diploid strains and then deriving haploid strains, it should be feasible to directly obtain promoter shut-off haploid strains by inserting an *enotetS* series promoter, along with the TetR-expressing cassette, upstream of the target essential gene using a single-step CRISPR-based marker-free knock-in (Jacobs et al., 2014; Zhang et al., 2018). The intended knock-in clones can be readily selected based on their ability to grow on ahTet-containing plates but not on ahTet-free plates. Promoter shut-off using the *nmt1* series promoters can be initiated by simply adding thiamine to the culture, whereas promoter shut-off using the *enotetS* series promoters requires the more laborious process of washing cells to remove ahTet. It would be worth exploring in the future the feasibility of implementing the method of light-induced inactivation of ahTet for shutting off the *enotetS* series promoters (Baumschlager et al., 2020).

## Materials and methods

### Plasmid construction

The plasmids used in this study are listed in Table S1, and the primers used in this study are listed in Table S2. We amplified the 276-bp *eno101* promoter (Wang et al., 2014) from genomic DNA using primers oYY1 and oYY2. This amplified promoter was then cloned between the SwaI and XhoI sites in the stable integration vector (SIV) pAV0714 (Addgene plasmid 133479), replacing the *act1* promoter (Vještica et al., 2020). The resulting plasmid was named pDB5530. For the construction of the *enotetS* promoter (*PenotetS*), a 19-bp sequence between the TATA box and the transcription start site (TSS) of the *eno101* promoter in pDB5530 was replaced with a 19-bp *tetO* sequence, specifically that of the O_2_ operator from the bacterial transposon Tn*10* (TCCCTATCAGTGATAGAGA) (Hillen et al., 1984, 1983; Wray and Reznikoff, 1983). This alteration was accomplished by performing overlapping PCR using pDB5530 as the template and oXH5, oXH6, oXH7, and oXH8 as primers, and then cloning the PCR product between the PacI and HpaI sites in pDB5530. The resulting plasmid was named pDB5531. To use *PenotetS* to drive the expression of the yellow fluorescent protein mECitrine, which serves as a reporter protein, we replaced the *adh1* promoter in pDB4915, a modified SIV plasmid containing mECitrine, with *PenotetS*. pDB4915 was derived from the SIV plasmid pAV0585 (pAde6^PmeI^-hphMX, Addgene plasmid 133472) (Vještica et al., 2020), by altering the linearization site from PmeI to NotI and introducing the *adh1* promoter, the coding sequence of mECitrine, and the terminator of the *Saccharomyces cerevisiae ADH1* gene. *PenotetS* was amplified from pDB5531 using primers oXH17 and oXH18, and then cloned between the ApaI and NheI sites in pDB4915. The resulting plasmid was named pDB5532.

To construct an integration plasmid that contains both *PenotetS* and a TetR-expressing cassette, we amplified a TetR expression cassette containing the cytomegalovirus promoter (*P.CMV*) and the coding sequence of TetR from pDM291-tetR-tup11Δ70 (Addgene plasmid 41027) (Zilio et al., 2012), using primers oXH19 and oXH47. The amplified cassette was then cloned into the KpnI site in pDB5532. The resulting plasmid was named pDB5318. To construct weakened variants of *PenotetS*, we used primers oXH34-oXH38 to amplify four PCR products, each containing *PenotetS* and a stem-loop-forming sequence immediately downstream of *PenotetS*. These PCR products were then cloned separately between the ApaI and NheI sites in pDB5318, generating four different plasmids, which were named pDB5319-pDB5322. The four weakened variants of *PenotetS* in these four plasmids were named *PenotetSW1*, *PenotetSW2*, *PenotetSW3*, and *PenotetSW4*. The plasmids pDB5318-pDB5322 have been deposited at Addgene (Addgene plasmids 204828-204832) and the Yeast Genetic Resource Center of Japan (YGRC/NBRP) (NBRP IDs FYP6242-6246). The SnapGene files for these plasmids are provided in a compressed folder as a supplementary file.

The DNA sequence encoding Mc-2A-Mi was synthesized by BGI Tech Solutions (Beijing Liuhe) and consists of three parts: the coding sequence of Mi, a 60-bp segment encoding the 20-amino acid T2A peptide (RAEGRGSLLTCGDVEENPGP) (Beekwilder et al., 2014), and the coding sequence of Mc. To construct a plasmid expressing Mc-2A-Mi, the Mc-2A-Mi sequence was amplified using primers oXH161 and oXH162, and then cloned between the NheI and BglII sites of pDB5319, replacing the coding sequence of mECitrine. In the resulting plasmid (pDB5533), Mc-2A-Mi is under the control of *PenotetSW1*.

The plasmids pDB4877, pDB4878, and pDB4879 were derived from the SIV vector pAV0585 (pAde6^PmeI^-hphMX, Addgene plasmid 133472) (Vještica et al., 2020), by altering the linearization site from PmeI to NotI and introducing the *nmt1* promoter, the *41nmt1* promoter, or the *81nmt1* promoter, respectively, along with the coding sequence of mECitrine, and the terminator of the *Saccharomyces cerevisiae ADH1* gene. The plasmids pDB5534, pDB5535, pDB5536, pDB5537, and pDB5538 were derived from the SIV plasmid pAV0586 (pLys3^BstZ17I^-bsdMX, Addgene plasmid 133473), by altering the linearization site from BstZ17I to NotI and introducing the TetR expression cassette, an *enotetS* series promoter, the coding sequence of mECitrine, and the terminator of the *Saccharomyces cerevisiae ADH1* gene. To construct plasmids expressing *nrs1*, the coding sequence of *nrs1* was amplified from genomic DNA and cloned into pDB5537 and pDB5536, replacing the coding sequence of mECitrine in each plasmid. The resulting plasmids were named pDB5539 and pDB5541, respectively. Two plasmids expressing *sui1* were similarly constructed, except cDNA was used as the amplification template, and they were named pDB5540 and pDB5542.

### Strain construction

The *S. pombe* strains used in this study are listed in Table S3. Strains containing an integrated plasmid were constructed by linearizing an SIV plasmid with NotI and then transforming the plasmid into a host strain. Correct integration of the plasmid into the *ade6* or *lys3* locus was verified using colony PCR (Vještica et al., 2020). Deletion strains were created using PCR-based gene deletion. Strains with a gene endogenously tagged at the C-terminus with the 3×sAID tag were generated through PCR-based gene tagging (Zhang et al., 2022). The *h+/h−* diploid strain DY51002 was constructed by crossing two haploid strains with *ade6-M210* and *ade6-M216* mutations, respectively. The two haploid strains were pre-grown on YES plates and then mixed in an equal ratio in 20 μL of water. The mixture was spotted on SPA plates with necessary supplements. After approximately 12 hours of incubation at 30°C, cells were collected and plated on YE plates containing 200 mg/L guanine at a density of approximately 10^4^ cells per plate. Guanine was used to inhibit the uptake of adenine, thereby preventing the growth of adenine auxotrophs and allowing only the growth of *ade6-M210*/*ade6-M216* diploid cells, which are adenine prototrophs due to interallelic complementation (Cummins and Mitchison, 1967; Smith, 2009). After incubating for 3 days at 30°C, colonies formed on the guanine-containing plates were streaked on YE2A plates. A diploid clone that showed high spore viability in tetrad analysis was saved as DY51002. A prototrophic *h+/h−* diploid strain, DY50999, constructed in the same way as DY51002, was plated on YEPD plates, and colonies that were unable to form spores were selected based on iodine staining. The mating type of the non-sporulating diploid clones was determined by colony PCR, and an *h+/h+* diploid strain was saved as DY51000.

### Media and growth conditions

The compositions of the media were as described (Petersen and Russell, 2016). Anhydrotetracycline hydrochloride (ahTet; J&K Scientific, catalog number 541642) was dissolved in DMSO at a concentration of 5 mg/mL and stored at −80°C.

To analyze the dose response of the inducer, the cells were first cultured in liquid YES medium or EMM medium with the necessary supplements until they reached the logarithmic phase. The DMSO-diluted ahTet stock was then added to achieve the desired concentrations, with an equal volume of DMSO serving as the control. After 24 hours of culturing (with cultures diluted when the OD600 approaching 1.0) at 30°C, logarithmic phase cells were collected for live-cell imaging.

For the time course assay of proliferating cells, the cells were first cultured in liquid YES medium or EMM medium with the necessary supplements until they reached the logarithmic phase at 30°C. After adding ahTet to the medium, the cells were collected at specific time points. During the time course, the cultures were diluted when the OD600 approached 1.0.

To quantify the expression levels of the *enotetS* series promoters, induced expression levels were measured after transferring logarithmic phase cells to media with 2.5 μg/mL ahTet for 24 hours at 30°C. Repressed expression levels were measured after transfer to media with DMSO under the same conditions.

To quantify the expression levels of the *nmt1* series promoters, the cells were first cultured in EMM medium containing thiamine until they reached the logarithmic phase. Then, to measure the induced expression levels, the cells were washed three times with water, transferred to EMM medium without thiamine, and grown at 30°C for 24 hours. To measure the repressed expression levels, the cells were cultured in EMM medium containing thiamine at 30°C for 24 hours.

For the time course assay of G0 phase cells, logarithmically growing cells grown in EMM medium were washed three times with water and shifted to EMM medium without nitrogen (EMM−N) for 24 hours at 30°C to arrest the cells in the G0 phase. After adding ahTet to the medium at a concentration of 2.5 μg/mL, the cells were collected at specific time points.

For the time course assay of spores, *h^90^* cells grown in YES medium were washed three times with water and shifted to MSL−N medium at 30°C for 48 hours to induce sporulation. Then, the cells were collected and treated with Snailase (Beijing Solarbio Science & Technology Co.,Ltd., catalog number S8280) in water at 30°C for 24 hours to digest non-spore cells. After digestion, spores were purified by centrifugation in Percoll solutions. Spores were incubated in EMM−N medium overnight. Then ahTet was added to the EMM−N medium. After adding ahTet to the medium at a concentration of 2.5 μg/mL, spores were collected at specific time points.

For the growth curve analysis, haploid deletion strains harboring a plasmid expressing the deleted gene from *PenotetSW2* or *PenotetSW3*, along with a wild-type strain were pre-cultured in YES liquid medium supplemented with ahTet until they reached the logarithmic phase at 30°C. Then, the cells were washed three times with water and transferred to either YES medium or YES medium supplemented with ahTet. The optical density at 600 nm (OD600) of each culture was measured at specific time points, with cultures being diluted if the OD600 approached 1.0. The number of generations was determined based on the OD600 and the dilution ratio.

### Fluorescence microscopy and image quantification

Live-cell imaging was performed using a Dragonfly 200 spinning disk confocal microscope system (Andor Technology) equipped with an mCherry/YFP/CFP filter set and an mCherry/GFP filter set. Images were acquired with a 100×, 1.4-NA objective using a Sona sCMOS camera and analyzed with Fiji (ImageJ).

To quantify the expression level of mECitrine, first, channels of images were split and the DIC channel was used to segment cell boundaries using Cellpose (version 2.1.1) (Stringer et al., 2021). The resulting masked images were processed with the LabelsToRois plugin (Waisman et al., 2021) in Fiji (Schindelin et al., 2012). This plugin generated outlines outlining the boundaries of each cell. Subsequently, the generated outlines were superimposed onto the mECitrine channel. Fiji was employed to calculate the mean fluorescence intensity for each individual cell.

In total, three images were acquired for each sample, with each image containing between 50 and 200 cells. The median fluorescence intensity of cells within each image was calculated and averaged across the triplicate images. Background fluorescence was measured using a parental strain that lacked mECitrine and was subsequently subtracted. The resulting fluorescence intensity provided a measurement of the expression level. For all fluorescence quantification results, we conducted two or more biological replicates, each yielding consistent outcomes.

### Induction and monitoring of meiosis

Log-phase cells grown in YES medium were collected, washed three times with water, resuspended in EMM medium without nitrogen (EMM−N), and incubated at 30°C for 24 hours to arrest cells in the G0 phase. Then, ahTet was added to the culture at a final concentration of 2.5 μg/mL to induce meiosis. At specific time points, cells were collected and fixed using 70% ethanol. The advancement of meiosis was evaluated by quantifying the number of nuclei per cell after staining the DNA with SYTOX Green (Thermo Fisher, catalog number S7020). For SYTOX Green staining, the fixed cells were washed once with water, resuspended in water containing 1 μM SYTOX Green, and incubated in the dark at room temperature for 5 minutes. The number of nuclei per cell was determined through imaging. The experiment was conducted in three biological replicates, with one replicate presented as representative data in the figure.

### Tetrad analysis

To examine spore viability of induced meiosis, cells collected 24 hours after meiosis induction were streaked onto YES plates. Tetrad analysis was carried out using a TDM50 tetrad dissection microscope (Micro Video Instruments, Avon, USA) on YES plates. To assess whether the exogenous expression of *nrs1* or *sui1* can complement the deletion of the corresponding endogenous gene, diploid strains freshly grown on YE2A plates were spotted onto SPASK plates. The SPASK plates were incubated at 30°C for about 24 h. Subsequently, tetrad analysis was carried out on YES plates with and without ahTet.

### Curve fitting

The induction kinetics data were fitted using Equation 1, as shown below.

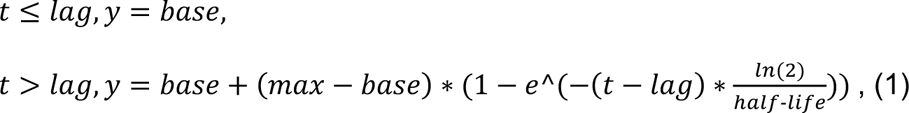

In this equation, “lag” represents the time required for the newly expressed reporter protein to become detectable, set to 0.5 hours because mECitrine, the reporter protein used, takes approximately 30 minutes to mature and emit fluorescence. “Base” denotes the expression level of the reporter protein at t = 0, set to the quantified value at that time. “Max” indicates the steady-state expression level. “Half-life” refers to the half-life of the reporter protein, determined by both protein degradation and dilution through cell doubling. Curve fitting was performed using the lmfit library in Python, employing the leastsq method (Newville et al., 2016).

### Disclosure on the use of artificial intelligence tools

The manuscript was proofread using the artificial intelligence tool editGPT. Following the use of this tool, the authors carefully reviewed and edited the content as necessary. The authors assume full responsibility for the publication’s content.

## Supporting information

Supplementary zip file containing SnapGene files

## Acknowledgements

We thank the Sophie Martin laboratory for making the SIV plasmids they have developed available through Addgene and the Yeast Genetic Resource Center of Japan (YGRC/NBRP). We thank YGRC/NBRP for providing the plasmids.

## Data and resource availability

Data availability: All relevant data can be found within the article and its supplementary information.

