## Supplementary Figures and Tables for "An improved tetracycline-inducible expression system for fission yeast"

**Figure S1.**

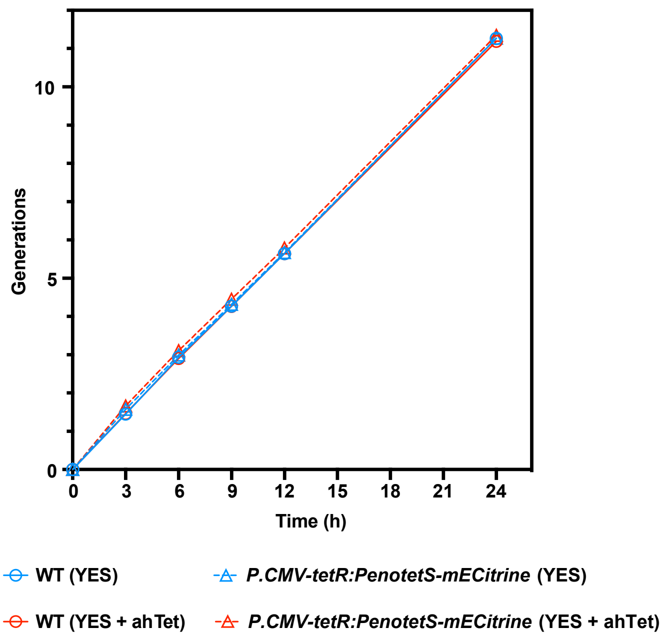


**Fig. S1. The presence of ahTet and the expression of TetR do not affect cell growth.**

Cells were first cultured in YES liquid medium without ahTet and then, at time 0, transferred to either ahTet-containing or ahTet-free YES media. The optical density at 600 nm (OD600) of each culture was measured at the specified time points. Cultures were diluted to maintain logarithmic growth when the OD600 approached 1.0. The number of generations was calculated using the OD600 measurements and dilution ratio.

### Figure S2.

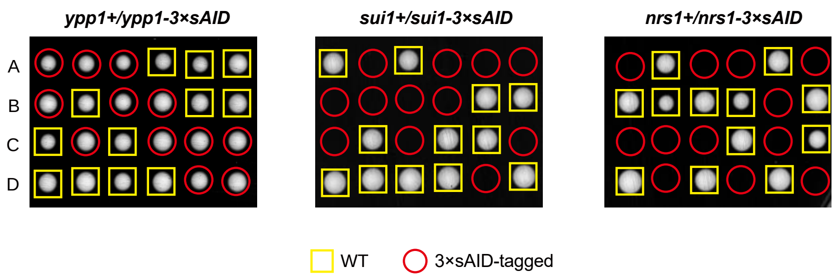


**Fig. S2. AID tagging of two essential genes caused lethality.**

Tetrad analysis was conducted on heterozygous diploids with one copy of an essential gene endogenously tagged with the 3×sAID tag (Zhang et al., 2022). Diploids were sporulated, and the resulting tetrads were dissected on YES plates.

**Table S1.** Plasmids used in this study

| Addgene ID | NBRP ID | Name | Descriptive name |
|---|---|---|---|
| | | **pDB5530** | pUra4-Peno276-sfGFP-Scer\T.CYC1 |
| | | **pDB5531** | pUra4-PenotetS-sfGFP-Scer\T.CYC1 |
| | | **pDB5532** | pAde6-PenotetS-mECitrine-Scer\T.ADH1:hphMX |
| 204828 | FYP6242 | **pDB5318** | pAde6-P.CMV-tetR:PenotetS-mECitrine-Scer\T.ADH1:hphMX |
| 204829 | FYP6243 | **pDB5319** | pAde6-P.CMV-tetR:PenotetSW1-mECitrine-Scer\T.ADH1:hphMX |
| 204830 | FYP6244 | **pDB5320** | pAde6-P.CMV-tetR:PenotetSW2-mECitrine-Scer\T.ADH1:hphMX |
| 204831 | FYP6245 | **pDB5321** | pAde6-P.CMV-tetR:PenotetSW3-mECitrine-Scer\T.ADH1:hphMX |
| 204832 | FYP6246 | **pDB5322** | pAde6-P.CMV-tetR:PenotetSW4-mECitrine-Scer\T.ADH1:hphMX |
| | | **pDB5533** | pAde6-P.CMV-tetR:PenotetSW1-McMi-Scer\T.ADH1:hphMX |
| | | **pDB3532** | pDM291-tetR-tup11Δ70 |
| | | **pDB4915** | pAde6-Padh1-mECitrine-Scer\T.ADH1:hphMX |
| | | **pDB4877** | pAde6-Pnmt1-mECitrine-Scer\T.ADH1:hphMX |
| | | **pDB4878** | pAde6-P41nmt1-mECitrine-Scer\T.ADH1:hphMX |
| | | **pDB4879** | pAde6-P81nmt1-mECitrine-Scer\T.ADH1:hphMX |
| | | **pDB5534** | pLys3-P.CMV-tetR:PenotetS-mECitrine-Scer\T.ADH1:bsdMX |
| | | **pDB5535** | pLys3-P.CMV-tetR:PenotetSW1-mECitrine-Scer\T.ADH1:bsdMX |
| | | **pDB5536** | pLys3-P.CMV-tetR:PenotetSW2-mECitrine-Scer\T.ADH1:bsdMX |
| | | **pDB5537** | pLys3-P.CMV-tetR:PenotetSW3-mECitrine-Scer\T.ADH1:bsdMX |
| | | **pDB5538** | pLys3-P.CMV-tetR:PenotetSW4-mECitrine-Scer\T.ADH1:bsdMX |
| | | **pDB5539** | pLys3-P.CMV-tetR:PenotetSW3-nrs1-Scer\T.ADH1:bsdMX |
| | | **pDB5540** | pLys3-P.CMV-tetR:PenotetSW3-sui1-Scer\T.ADH1:bsdMX |
| | | **pDB5541** | pLys3-P.CMV-tetR:PenotetSW2-nrs1-Scer\T.ADH1:bsdMX |
| | | **pDB5542** | pLys3-P.CMV-tetR:PenotetSW2-sui1-Scer\T.ADH1:bsdMX |

**Table S2.** Primers used in this study

| Primer | Sequence |
|---|---|
| **oYY1** | GGTACCGGGCCCATTTAAATCATCTCTTGCCCCTT |
| **oYY2** | AAGAATTCGTCGACCTCGAGCGATGTTTACTGTAGAATAC |
| **oXH5** | TACCCTAACGTTCCGGTTAACGATT |
| **oXH6** | TCTCTATCACTGATAGGGACACTCTATATATACCTGGAGGAAGC |
| **oXH7** | GTCCCTATCAGTGATAGAGACTTGTTTCAGTAAGAATCAATTAGTATTCT |
| **oXH8** | TTGGAGCTTGCCATGTTAATTAAGA |
| **oXH17** | GGCGAATTGGGTACCGGGCCCCATCTCTTGCCCCTTCTAAG |
| **oXH18** | TTCTCCTTTACTCATGCTAGCCGATGTTTACTGTAGAATACTAATTGATT |
| **oXH19** | GGGGCAAGAGATGGGGCCCGGGGGGTCGAGGAGCTT |
| **oXH47** | ACTCACTATAGGGCGAATTGGTTAAGATCCACTTTCACATTTAAGTTG |
| **oXH34** | GCCAAGCTCCTCGACCCCCCGGGCCCCATCTCTTGCCCCTTCTAAG |
| **oXH35** | AGTTCTTCTCCTTTACTCATGCTAGCGCCCGAGCGGCCGATGTTTACTGTAG AATACTAATTGATTC |
| **oXH36** | AGTTCTTCTCCTTTACTCATGCTAGCGGCCCGAGCGGCCCGATGTTTACTG TAGAATACTAATTGATTC |
| **oXH37** | AGTTCTTCTCCTTTACTCATGCTAGCGGGCCCGAGCGGCCCCGATGTTTAC TGTAGAATACTAATTGATTC |
| **oXH38** | AGTTCTTCTCCTTTACTCATGCTAGCCGGGCCCGAGCGGCCCGCGATGTTT ACTGTAGAATACTAATTGATTC |
| **oXH161** | ATCGGCCGCTCGGGCGCTAGCATGTCAGCAGAAGACCTTTT |
| **oXH162** | AGAAGTGGCGCGTTAAGATCTTTATCTTTTAACTTTATTCTTTCTAGGCTGG |
| **oXH163** | CGGGCCGCTCGGGCCGCTAGCATGTCAGCAGAAGACCTTTT |
| **oXH164** | GGGCCGCTCGGGCCCGCTAGCATGTCAGCAGAAGACCTTTT |
| **oXH165** | GGCCGCTCGGGCCCGGCTAGCATGTCAGCAGAAGACCTTTT |

**Table S3.** Strains used in this study

| Name | Alias | Mating Type | Genotype | Use |
|---|---|---|---|---|
| DY49190 | LD1 | *h−* | *leu1-32 ura4-D18* | Fig. 1D,2B,2D,3B,3C,3D,3E,4 |
| DY50985 | LV91 | *h−* | *leu1-32 ura4-D18 ade6+::P.CMV-tetR:PenotetS-mECitrine-Scer\T.ADH1:hphMX* | Fig. 1D,3A,3B,3C,4 |
| DY50986 | LV93 | *h−* | *leu1-32 ura4-D18 ade6+::P.CMV-tetR:PenotetS-mECitrine-Scer\T.ADH1:hphMX* | Fig. 2 |
| DY50987 | LV171 | *h−* | *leu1-32 ura4-D18 ade6+::P.CMV-tetR:PenotetSW1-mECitrine-Scer\T.ADH1:hphMX* | Fig. 1D,4 |
| DY50988 | LV173 | *h−* | *leu1-32 ura4-D18 ade6+::P.CMV-tetR:PenotetSW2-mECitrine-Scer\T.ADH1:hphMX* | Fig. 1D,3D,3E,4 |
| DY50989 | LV175 | *h−* | *leu1-32 ura4-D18 ade6+::P.CMV-tetR:PenotetSW3-mECitrine-Scer\T.ADH1:hphMX* | Fig. 1D,4 |
| DY50990 | LV177 | *h−* | *leu1-32 ura4-D18 ade6+::P.CMV-tetR:PenotetSW4-mECitrine-Scer\T.ADH1:hphMX* | Fig. 1D,4 |
| DY50994 | LV70 | *h−* | *leu1-32 ura4-D18 ade6+::Padh1-mECitrine-Scer\T.ADH1:hphMX* | Fig. 1D,4A |
| DY50991 | LV61 | *h−* | *leu1-32 ura4-D18 ade6+::Pnmt1-mECitrine-Scer\T.ADH1:hphMX* | Fig. 4B,4C |
| DY50992 | LV64 | *h−* | *leu1-32 ura4-D18 ade6+::P41nmt1-mECitrine-Scer\T.ADH1:hphMX* | Fig. 4B,4C |
| DY50993 | LV67 | *h−* | *leu1-32 ura4-D18 ade6+::P81nmt1-mECitrine-Scer\T.ADH1:hphMX* | Fig. 4B,4C |
| DY49197 | LD331 | *h+* | | Fig. 5C,7B,S1 |
| DY47073 | DY47073 | *h90* | | Fig. 5C |
| DY50995 | LV598 | *h+* | *ade6+::P.CMV-tetR:PenotetS-mECitrine-Scer\T.ADH1:hphMX* | Fig. 5C, S1 |
| DY50996 | LV599 | *h+* | *ade6+::P.CMV-tetR:PenotetS-mECitrine-Scer\T.ADH1:hphMX* | Fig. 5A,5C |
| DY50998 | LV603 | *h90* | *ade6+::P.CMV-tetR:PenotetS-mECitrine-Scer\T.ADH1:hphMX* | Fig. 5B,5C |

| | | | | |
|---|---|---|---|---|
| DY50999 | LV868 | *h+/h−* | *mat1M-Δ17/mat1P-Δ17 ade6-M216/ade6-M210 leu1+/leu1-32* | |
| DY51000 | LV863 | *h+/h+* | *mat1P-Δ17/mat1P-Δ17 ade6-M216/ade6-M210 leu1+/leu1-32* | |
| DY51001 | LV882 | *h+/h+* | *mat1P-Δ17/mat1P-Δ17 leu1+/leu1-32 ade6-M216/ade6-M210 ade6::P.CMV-tetR:PenotetSW1-McMi-Scer\T.ADH1:hphMX* | Fig. 6 |
| DY51002 | LV267 | *h+/h−* | *mat1M-Δ17/mat1P-Δ17 ade6-M210/ade6-M216 leu1-32/leu1-32* | |
| DY51003 | LV680 | *h+/h−* | *mat1M-Δ17/mat1P-Δ17 leu1-32/leu1-32 ade6-M210/ade6-M216 sui1+/sui1Δ::NATsNG* | |
| DY51004 | LV488 | *h+/h−* | *mat1M-Δ17/mat1P-Δ17 leu1-32/leu1-32 ade6-M210/ade6-M216 nrs1+/nrs1Δ::NATsNG* | |
| DY51005 | LV1047 | *h+/h−* | *mat1M-Δ17/mat1P-Δ17 leu1-32/leu1-32 ade6-M210/ade6-M216 sui1+/sui1Δ::NATsNG lys3+/lys3+::P.CMV-tetR:PenotetSW3-sui1-Scer\T.ADH1:bsdMX* | Fig. 7A |
| DY51006 | LV1051 | *h+/h−* | *mat1M-Δ17/mat1P-Δ17 leu1-32/leu1-32 ade6-M210/ade6-M216 sui1+/sui1Δ::NATsNG lys3+/lys3+::P.CMV-tetR:PenotetSW2-sui1-Scer\T.ADH1:bsdMX* | Fig. 7A |
| DY51007 | LV1042 | *h+/h−* | *mat1M-Δ17/mat1P-Δ17 leu1-32/leu1-32 ade6-M210/ade6-M216 nrs1+/nrs1Δ::NATsNG lys3+/lys3+::P.CMV-tetR:PenotetSW3-nrs1-Scer\T.ADH1:bsdMX* | Fig. 7A |
| DY51008 | LV1045 | *h+/h−* | *mat1M-Δ17/mat1P-Δ17 leu1-32/leu1-32 ade6-M210/ade6-M216 nrs1+/nrs1Δ::NATsNG lys3+/lys3+::P.CMV-tetR:PenotetSW2-nrs1-Scer\T.ADH1:bsdMX* | Fig. 7A |
| DY51009 | LV1071 | ? | *mat1M-Δ17(or mat1P-Δ17) leu1-32 ade6-M210(or ade6-M216) nrs1Δ::NATsNG lys3+::P.CMV-tetR:PenotetSW3-sui1-Scer\T.ADH1:bsdMX* | Fig. 7B |
| DY51010 | LV1074 | ? | *mat1M-Δ17(or mat1P-Δ17) leu1-32 ade6-M210(or ade6-M216) nrs1Δ::NATsNG lys3+::P.CMV-tetR:PenotetSW2-sui1-Scer\T.ADH1:bsdMX* | Fig. 7B |
| DY51011 | LV1068 | ? | *mat1M-Δ17(or mat1P-Δ17) leu1-32 ade6-M210(or ade6-M216) nrs1Δ::NATsNG lys3+::P.CMV-tetR:PenotetSW3-nrs1-Scer\T.ADH1:bsdMX* | Fig. 7B |

| | | | | |
|---|---|---|---|---|
| DY51012 | LV1070 | ? | *mat1M-Δ17(or mat1P-Δ17) leu1-32 ade6-M210(or ade6-M216) nrs1Δ::NATsNG lys3+::P.CMV-tetR:PenotetSW2-nrs1-Scer\T.ADH1:bsdMX* | Fig. 7B |
| DY51013 | ZXRY4876 | *h+/h−* | *mat1P-Δ17/mat1M-Δ17 leu1-32/leu1-32 ade6-M210/ade6-M216 lys3+/lys3+::Padh1-OsTIR1-F74A:bsdMX* | |
| DY51014 | LV684 | *h+/h−* | *mat1P-Δ17/mat1M-Δ17 leu1-32/leu1-32 ade6-M210/ade6-M216 lys3+/lys3+::Padh1-OsTIR1-F74A:bsdMX ypp1+/ypp1-3xsAID:kanMX* | Fig. S2 |
| DY51015 | LV702 | *h+/h−* | *mat1P-Δ17/mat1M-Δ17 leu1-32/leu1-32 ade6-M210/ade6-M216 lys3+/lys3::Padh1-OsTIR1-F74A:bsdMX sui1+/sui1-3xsAID:kanMX* | Fig. S2 |
| DY51016 | LV694 | *h+/h−* | *mat1P-Δ17/mat1M-Δ17 leu1-32/leu1-32 ade6-M210/ade6-M216 lys3+/lys3::Padh1-OsTIR1-F74A:bsdMX nrs1+/nrs1-3xsAID:kanMX* | Fig. S2 |
